# A portable and wind resistant drift fence array for arid environments

**DOI:** 10.1101/2023.09.28.560034

**Authors:** Benjamin T. Camper, Zachary T. Laughlin, Andrew S. Kanes, Riley T. Manuel, Sharon A. Bewick

## Abstract

Drift fences are passive trapping systems for both small vertebrates and large invertebrates. Most drift fence designs are semi-permanent or otherwise difficult to transport after initial installation. While these designs are effective for replicate trapping through time, most designs lack portability. Here, we propose a novel drift fence design that uses PVC pipes and fiberglass mesh screen. The combination of hollow PVC pipes and mesh screen creates a lightweight system that facilitates rapid deployment and redeployment across locations. Since the PVC pipes can be filled with topsoil from the site of trap installation, enhanced portability does not come at the cost of fence stability or wind resistance. We provide results on trap performance in a New Mexico flatland desert and discuss the efficacy and cost of our proposed drift fence design.

## Introduction

Drift fences paired with traps (henceforth drift fence arrays) are an effective method to monitor terrestrial and aquatic fauna (e.g., reptiles, amphibians, small mammals, invertebrates, etc.), both through time and across habitats (Ali et al., 2018; Brennan et al., 2005; Maritz et al., 2007; Murphy et al., 2007; Willson & Dorcas, 2004). These traps work by intercepting small animals as they traverse the environment. When an animal meets a drift fence bisecting its path, the animal will typically turn and continue along the fence where it is then captured by traps placed on either side or at the centre (but see Amber et al. [2021] for an introduction to camera trap designs). Drift fences are advantageous because they are passive monitoring efforts. This is particularly useful for mobile terrestrial and aquatic organisms that otherwise evade capture due to their cryptic lifestyle, rarity, or escape behaviour. Drift fences are also durable, which enables replicate sampling (i.e., installing fencing and then leaving it semi-permanently). However, drift fence durability and permanence typically come at the cost of portability. As a result, drift fences are less useful in study designs where fences must be repeatedly uninstalled at certain sites and reinstalled at others. This is a common practice during site selection or in situations where only a handful of animals are required from any given location. Consequently, it is impractical to use most drift fence designs except for long-term studies. This is especially true if drift fences are accompanied by semi-permanent traps such as pitfall traps.

While most drift fence arrays follow the same general design and operate based on the same general principals, array shape and size (Brennan et al., 2005; Ellis, 2013), array material (Enge, 1997), trap shape and size (Maritz et al., 2007; Todd et al., 2007), and trap type (Greenberg et al., 1994) must all be taken into consideration. These aspects of design are typically chosen based on the broader study goals. Factors to consider include the target habitat, the species of interest, and material costs (Enge, 1997; Todd et al., 2007). For example, when monitoring terrestrial amphibians during spring migration to breeding ponds, it is common to use a linear fence design that encircles the wetland perimeter (Semlitsch, 1983). In these instances, porous materials such as silt fencing and funnel traps are often preferable, as this prevents inundation and trap displacement during periods of wetland flooding (Enge, 1998). By contrast, when monitoring active foraging reptiles that shuttle between desert microclimates to regulate body temperature (Bowker et al., 1986), there may be no *a priori* assumptions about movement pattern directionality. For this type of scenario, a four– (‘X’-) or three-fence (‘Y’-shaped) fence design may be implemented to capture reptile movement spanning 360° from the fence centre (Ellis, 2013). Further, while flooding may be less relevant in desert environments, stability under high wind conditions may be paramount.

Some aspects of drift fence array design have been better explored than others. Numerous studies have examined the influence of drift fence array shape (Brennan et al., 2005; Ellis, 2013; Maritz et al., 2007; Todd et al., 2007). Ellis (2013), for example, found that two-fence (‘X’-shaped) trapping arrays out-performed one-fence (‘I’-shaped) arrays when comparing reptile and small mammal capture rates in the floodplains of the Southern Mallee Region of New South Wales, Australia. While this is broadly consistent with findings from a similar study by Morton et al. (1988), Hobbs et al. (1994) found that one-fence designs are superior in the same region of Australia. Thus, even in similar habitats, system-specific or researcher effects can drive substantial variation in the efficacy of a drift fence array design. In addition to fence shape, fence materials have also been extensively explored. Common options include silt fencing or metal flashing, paired with wooden or metal rebar stakes. Whereas the water permeability and lightweight nature of silt fencing can be a benefit in wet conditions or with animals not prone to climbing, the permanence and sleekness of metal flashing can be advantageous in windy conditions or where animals are more likely to scale the fence (Enge, 1997, 1998). Other, more subtle aspects of fence design have been studied as well. Ellis (2013), for example, demonstrated differences between opaque ‘dampcourse’ fencing and translucent ‘flyscreen’ fencing, with reptiles being more frequently captured by the dampcourse fencing while small mammals were more frequently captured by the flyscreen fencing. Beyond fence shape and material, trap choice has also received a fair amount of attention. For instance, Todd et al. (2007) compared the effect of trap size on herpetofaunal capture rates in a Carolina bay (South Carolina, USA) and found that small pitfall traps were largely ineffective compared to large pitfall traps. Todd et al. (2007) also considered the impact of trap type – including pitfall traps (Todd et al., 2007) and funnel traps – across different animal groups (Ali et al., 2018; Camper, 2005). Interestingly, larger pitfall traps were more effective at capturing species of turtles, small snakes, salamanders, and toads, whereas box funnel traps were most effective at capturing species of lizards, large snakes, and other frogs.

One aspect of drift fence array design that has received less attention is portability. Although funnel traps are commonly implemented as portable alternatives to pitfall traps for flexible trapping schemes, few alternatives have been offered for the fencing itself. In particular, there are not many alternatives to burying fencing and staking posts into the ground, which is a labour-intensive procedure. One exception is Rice et al. (2006), who proposed a portable fence design for herpetofaunal surveys in the Everglades National Park. While the Rice et al. (2006) design worked well in the wetland habitat of the everglades, it is less appropriate for conditions that require durability and stability under high wind conditions. No studies, to our knowledge, have addressed balancing the trade-off between wind resistance and portability. Our goal was to develop a highly portable, but wind-stable drift fence array that maximized capture rates of lizards from flatland desert ecosystems across the Rio Grande Valley in New Mexico, USA. Specifically, we were targeting whiptail lizards (genus *Aspidoscelis*). These are highly mobile reptiles with a tendency to escape down nearby burrows or into nearby shrubs (Schall & Pianka, 1980). As a result, desert *Aspidoscelis* species are challenging to capture via active methods (hand-capture, noosing, rubber band stunning, etc.). This makes passive capture efforts, such as drift fence arrays, the best option. Our study sites spanned a >150 km range across Socorro and Doña Ana counties, stretching from central to southern New Mexico. Despite the wide geography that we covered, our sites were generally characterized by flat terrain, sandy soil, and grass– or shrub-dominated plant communities with frequent high wind speeds upwards of 14 m/s. Additionally, New Mexico’s Rio Grande Valley harbours many important archaeological sites (Shackley, 2013; Shackley & Huckell, 2020). Thus, for numerous land stewards, archaeological disturbance is a point of concern, and any need for extensive ground disturbance can limit or fully restrict activities in certain locations.

Given the goals of our study, the traits of our study organisms, regional characteristics, and site restrictions, we designed our drift fence arrays with the following target specifications:

(1) manipulation by limited personnel (sometimes by one person),
(2) portability (easy redeployment across sites),
(3) wind resistance (the ability to resistant wind speeds frequenting 12–14 m/s) (Zlotin & Parmenter, 2008),
(4) limited ground disturbance (reducing environmental damage, and limiting the probability of disturbing archaeological remains).

Our novel drift fence array design uses hollow, soil, or sand-filled PVC pipes and fiberglass mesh screen (commonly used to screen porches). This lightweight construction, paired with hollow PVC pipes that can be weighed down with *in-situ* materials achieves portability requirements, while still maintaining stability under high wind conditions as well as high lizard capture rates. Further, in contrast to common drift fence methods, our design offers less ground disturbance because we do not bury the fencing nor stake the posts. Our design may aid those seeking a balance between portability and wind resistance, as well as those wishing to mitigate ground disturbance. These can be important considerations for people performing fieldwork in arid ecosystems. This is particularly true for researchers who wish to select trapping locations more haphazardly, for example, when the target species are stochastically distributed or exhibit some temporal trend in location, or when collecting genetic samples where only a small sample size is necessary at a particular location.

## Methods

### Materials and Assembly

The drift fence for each trapping array was constructed using the following materials: 24 m [26.25 ft] of fiberglass mesh screen (height 91.44 cm [3 ft]; 2.071 kg [4.622 lb]; $36.34), 5.69 m [18.668 ft] of PVC pipe (diameter 5.08 cm [2 in]; 3.19 kg [7.037 lb]; $37.34), four five-way PVC connectors (diameter 5.08 cm [2 in]; 0.816 kg [1.8 lb]; $39.18), 16 PVC caps (diameter 5.08 cm [2 in]; 2.032 kg [4.48 lb]; $28.80), 16 nylon cable ties (length 27.94 cm [11 in] **×** width 0.508 cm [0.2 in]; 33.152 g [1.184 oz]; $1.44), 10.973 m [12 ft] of duct tape (45.36 g [1.62 oz]; $1.59), and 21 tent stakes (length 17.78 cm [7 in]; 0.543 kg [1.196 lb]; $14.70). At locations with high wind speeds (upwards of 14 m/s), we also used woven polypropylene sandbags (length 35.56 cm [14 in] **×** height 66.04 cm [26 in]) to reinforce each fence corner. For each trapping array, we lined the drift fence with 6 double-sided rectangular prawn (i.e., box funnel) traps (brand KUFA sports; model S44X2N; each length 45.72 cm [18 in] **×** height 20.32 cm [8 in] **×** width 30.48 cm [12 in]; each 0.851 g [1.9 lb]; each $25) topped with wooden cover boards (ca. length 60.96 cm [24 in] **×** height 0.9 cm [0.35 in] **×** width 45.72 cm [18 in]; 0.680 kg [1.5 lb]; ca. $5) at the midpoint of each fence. We purchased our materials for fiberglass mesh screen, five-way PVC connectors, PVC caps, nylon cable ties, and duct tape from the most inexpensive vendors on March, 2022 from Amazon.com. We also purchased prawn traps from Amazon.com, but we strictly filtered for rectangular designs, enhanced sturdiness, and double-sidedness. We purchased PVC pipe and cover boards at Lowe’s Home Improvement and tent stakes from Walmart. The total cost and weight of a single drift fence array – excluding funnel traps, shade boards, and sand bags – was ca. $159.39 and 7.943 kg, respectively.

To construct a trapping array, we cut the fiberglass mesh screen (hereafter “screen”) into three 8 m **×** 91.44 cm [26.25 **×** 3 ft] lengths, each representing a fence ‘segment’. We then reinforced each 91.44 cm [3 ft] edge with two 91.44 cm [3 ft] lengths of duct tape, one for each side of the screen edge. Next, we cut the 5.08 cm [2 in] diameter PVC pipe into four 60.96 cm [24 in] lengths and sixteen 20.32 cm [8 in] lengths using PVC cutters. We then cable-tied separate ‘outer’ 60.96 cm [24 in] PVC posts to one end of each screen segment and a common ‘central’ 60.96 cm [24 in] PVC post to the other end of each screen segment. This resulted in a triangular ‘Y’ shape (see Figure 1) inspired by Jones (1986) and K. M. Enge (1998) where each screen segment serves as a separate fence, with three PVC posts at the three ends of the fences and a single PVC post at the centre. Note that the screen height was 91.44 cm [3 ft] and the PVC post heights were 60.96 cm [24 in]. We matched the top of one PVC post and the screen before cable-tying, leaving 60.96 cm [24 in] of screen unattached. We folded this extra 30.48 cm [12 in] portion upon itself 3–5 times, until it was 8–20 cm in length. We then staked the excess screen to the ground in the field (see description below). Finally, to construct PVC post bases (anchoring the posts to the ground), we connected four 20.32 cm [8 in] PVC pipes to a five-way PVC connector and attached PVC caps to each segments’ end.

**Figure 1.**
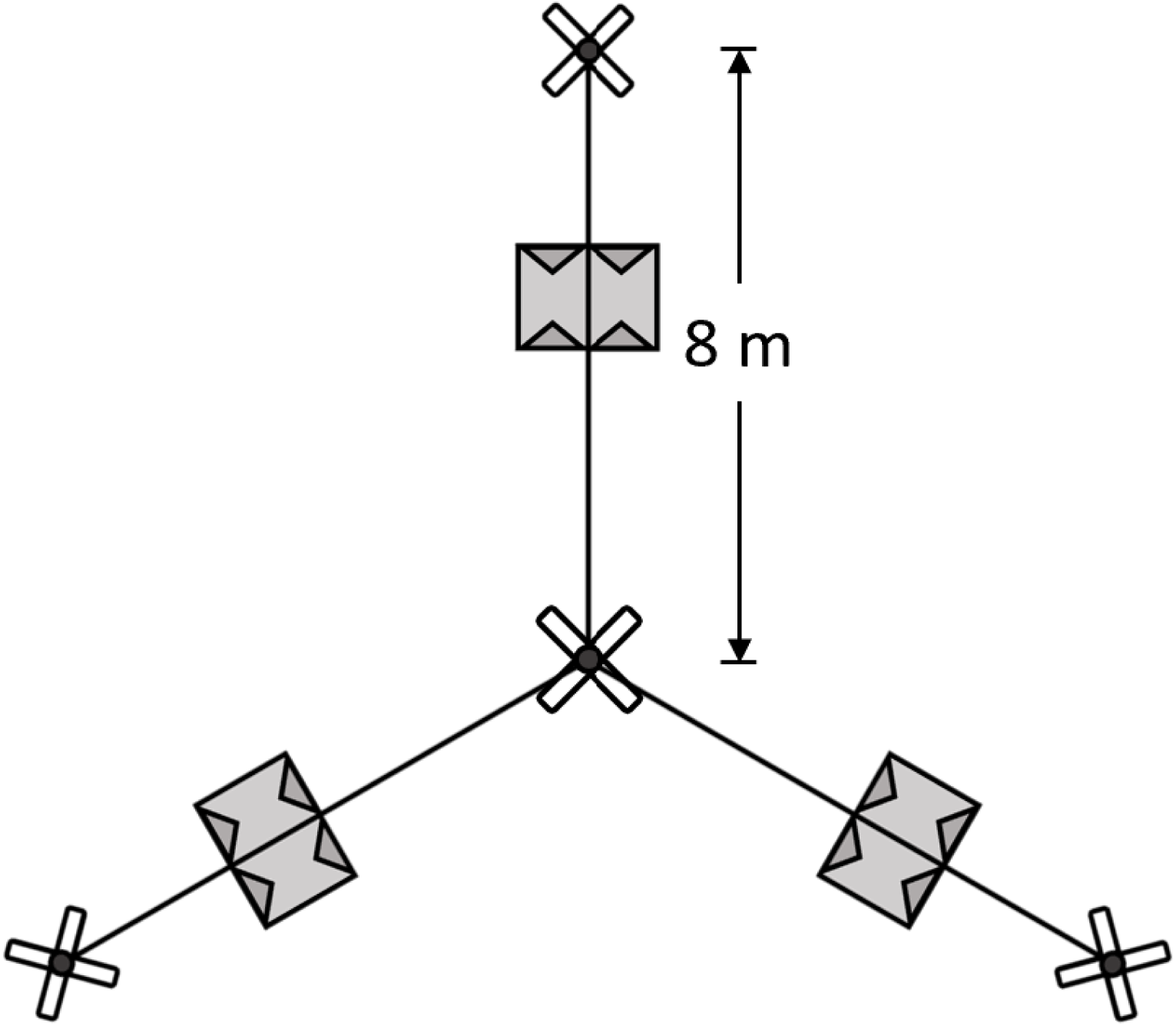
Top-down view of drift fence array configuration. Each fence is 8 m in length. Box funnel traps are depicted as grey rectangles, and PVC post bases, at each fence fence’s extremity, are in white.

### Field Assembly

PVC posts attached to screen were rolled into bundles for transportation to and from the field and in between field sites (see Table 1 for site details). We erected each trapping array by unrolling the fence bundles and connecting the PVC bases to the PVC posts by attaching the fifth empty connector socket to the PVC post. Before connecting the PVC base to the PVC post, we filled the base with loose topsoil (i.e., sand). In areas where this could be problematic due to ground disturbance restrictions, bringing heavy metal chains (Rice et al., 2006) or additional bags of sand are viable alternatives, though less portable and more expensive. After connecting the base to the post, we filled the post with topsoil to again serve as a weighted anchor for each fence segment. Following assembly, we positioned our drift fence in a Y-shape for 360° sampling by stretching each 8 m fence 120° from the adjacent fence (Figure 1). Lastly, any necessary vegetation was cleared, the dangling 30.48 cm [12 in] screen flap at the bottom of the fence was folded upon itself 3–5 times to make it flush with the ground, and seven tent stakes were pounded through the folded screen and into the ground (see Figure 2 for an example in the field). In instances where the folded fences were not completely flush with the ground, we brushed loose topsoil over this folded section.

**Figure 2.**
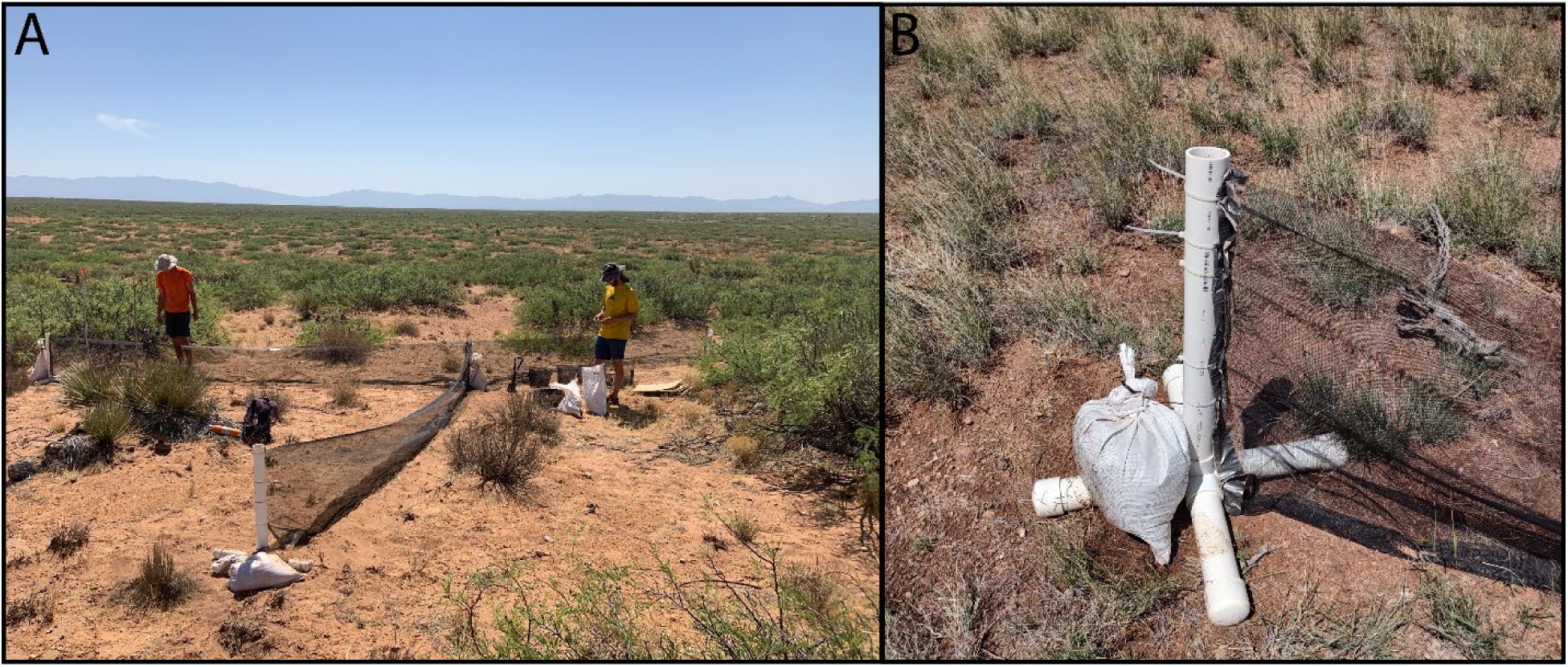
Trapping array design in the field. A) Installed trapping array including drift fence, box funnel traps, and shade cover boards. B) Close-up depiction of PVC post and bases. Photos (A) by Benjamin T. Camper and Andrew S. Kanes (B).

**Table 1.** Overview of summer 2022 trapping results and sampling effort with our novel drift fence trapping arrays. Total trapping days (n^TD^) and total trap checks (n^TC^) are reported as the frequency of days and checks respectively in which reptiles were present. Note that sparse forbes and grasses were found at each site. GPS coordinates are in Decimal Degrees (WGS84). Note for site names, ‘BLM’ denotes Bureau of Land Management ownership, ‘S’ denotes Sevilleta National Wildlife Refuge ownership, and ‘J’ denotes USDA Jornada Rangeland ownership. For Jornada Rangeland sites, ‘M’ and ‘MS’ indicate the vegetation distinction in the site description (Site^D^).

| Site | County | $n^R$ | $n^A$ | $n^{TD}$ | $n^{TC}$ | Site<sup>D</sup> | Lat | Long |
| --- | --- | --- | --- | --- | --- | --- | --- | --- |
| BLM-NW | Socorro | 6 | 66 | 10 | 18 | Grassland with sparse creosote shrub | 34.442710 | -106.922640 |
| S-NW | Socorro | 9 | 128 | 26 | 49 | Heterogeneous grassland-mesquite shrubland | 34.397882 | -106.867637 |
| S-NE | Socorro | 6 | 41 | 11 | 21 | Grassland with sparse cholla cactus | 34.331178 | -106.636717 |
| J-M | Doña Ana | 3 | 59 | 9 | 18 | Mesquite shrubland | 32.603865 | -106.852205 |
| J-MS | Doña Ana | 8 | 186 | 22 | 45 | Mesquite-sagebrush shrubland | 32.610940 | -106.800121 |
| **Total:** |  | 15 | 480 | 78 | 151 |  |  |  |

Trapping arrays were carried between 100 m and 500 m from the road for trapping. We found that two to three people could carry all the materials – fencing, bases, trowels, mallet, sand bags, box funnel traps, and shade cover boards – for a single trapping array in one trip (although this could vary depending on the physical ability of the field team). Construction time (not including material transportation) took between 30 and 45 minutes between two people. However, installation time varied depending on how much vegetation needed to be disturbed and how level the ground was. Array deconstruction was generally much faster, at about 20-30 minutes between two people.

### Field Survey

We used our novel trapping arrays to perform a summer-long survey of reptiles (Appendix 1) across 5 sites in central New Mexico (Table 1) between May-August 2022. Our sampling effort varied through time and by site depending on capture rates of our focal *Aspidoscelis* spp. Once deployed at a particular site, traps were operated continuously, except at high wind speeds (ca. 12–14 m/s). During times of high wind, we would pause our trapping endeavours. This was done due to low lizard activity (personal observation, June 15, 2019) on windy days, rather than any issue with trap operation under such conditions. In general, six trapping arrays were placed at each site and checked thrice daily to mitigate lizard stress and prevent desiccation. We recorded reptile presences across drift fence arrays, sites, dates, and times.

### Trap Comparison

At one site that we surveyed in 2022 – ‘Sevilleta Northeast’ in Socorro County (see Table 1 for more information) – we had previously performed a similar survey effort with standard silt fencing in 2021. We used this to provide a preliminary comparison between the performance of our new trapping arrays and the silt fencing trapping arrays. The silt fencing arrays had four 9.144 m [30 ft] fences in comparison to three 8 m fences in our new design. Like our new design, each silt fence maintained two box funnel traps.

Using the R statistical software (version 4.2.1; R Core Team, 2022, we compared these two drift fence array types across 11 days spanning late May to early June. We used a nonparametric Kruskal-Wallis Test (function ‘*kruskal.test*’) to test for differences in species richness and abundance of reptiles across array types. To account for the effect of the differing numbers of traps and fence lengths between array designs (see Table 2), we ran multiple Kruskal-Wallis tests controlling for these design differences by dividing abundance and species richness values by the number of traps and length of fencing.

**Table 2.**
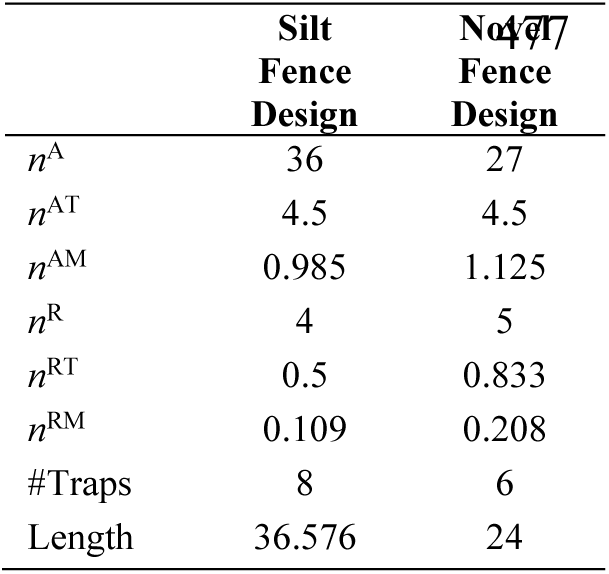
Summary results for total abundance and species richness of reptiles between a silt fence drift fence array design and our novel design at the ‘Sevilleta Northeast’ site for 11 days. Total abundance (’^A^’) and species richness (’^R^’) values indicated by ‘^T^’ indicate total abundance or species richness values divided by the number of traps used in either design. Abundance and species richness values indicated by ‘^M^’ indicate total abundance or species richness values divided by the length of fencing (meters) used in either design.

We also used the ‘*adonis2’* function (PERMANOVA) from the ‘vegan’ R package (version 2.6-2; Oksanen et al., 2013) to test for differences in community composition of reptiles across drift fence array types. For this approach, we removed the effect of the differing number of traps and fence lengths between array designs by rescaling our community matrix to reflect proportional abundance.

Code and data relevant to this manuscript are made available at https://github.com/camperbencamper/NM_portable_trapping_array.

## Results

### Survey Results

Using our novel drift fence array, we captured 480 reptiles of 15 species across 5 sites (Appendix 1) distributed between Socorro and Doña Ana counties, New Mexico in summer 2022. We asymmetrically sampled sites based off rates of capture of different species of whiptail lizards. For this reason, we report presence-only sampling effort for our broader community study (Table 1) for both sampling days (78 total) and sampling trap checks (151 total).

### Comparing Array Designs

In our comparison between drift fence array designs, based on 11 comparable days in May-June at Sevilleta Northeast, we captured 36 reptiles of four species (*Aspidoscelis inornatus*, *Aspidoscelis neomexicanus*, *Holbrookia maculata*, and *Plestiodon obsoletus*) and 27 reptiles of five species (*A. inornatus*, *A. neomexicanus*, *Coluber flagellum*, *H. maculata*, and *Pituophis catenifer*) for the silt fencing array design and our novel array design respectively (Table 2). Although this is a higher total capture rate for the silt fence array design, the novel array design outperformed the silt fence array design when controlling for the number of traps and the lengths of fencing respectively (see Table 2).

To determine whether differences in capture rates between the two designs were statistically significant, we aggregated reptile sampling by date. We then used a Kruskal-Wallis Test to test for the effect of drift fence array design on total reptile abundance and reptile species richness (see Table 3 & Figure 3). We found that differences in reptile abundance between the two array designs were not statistically significant (p-value=0.2771), but that differences in reptile species richness were statistically significant (p-value=0.04444). When controlling for the number of traps and length of fencing for each array design, however, neither differences in reptile species richness nor differences in reptile abundance were statistically significant (see Table 3).

**Figure 3.**
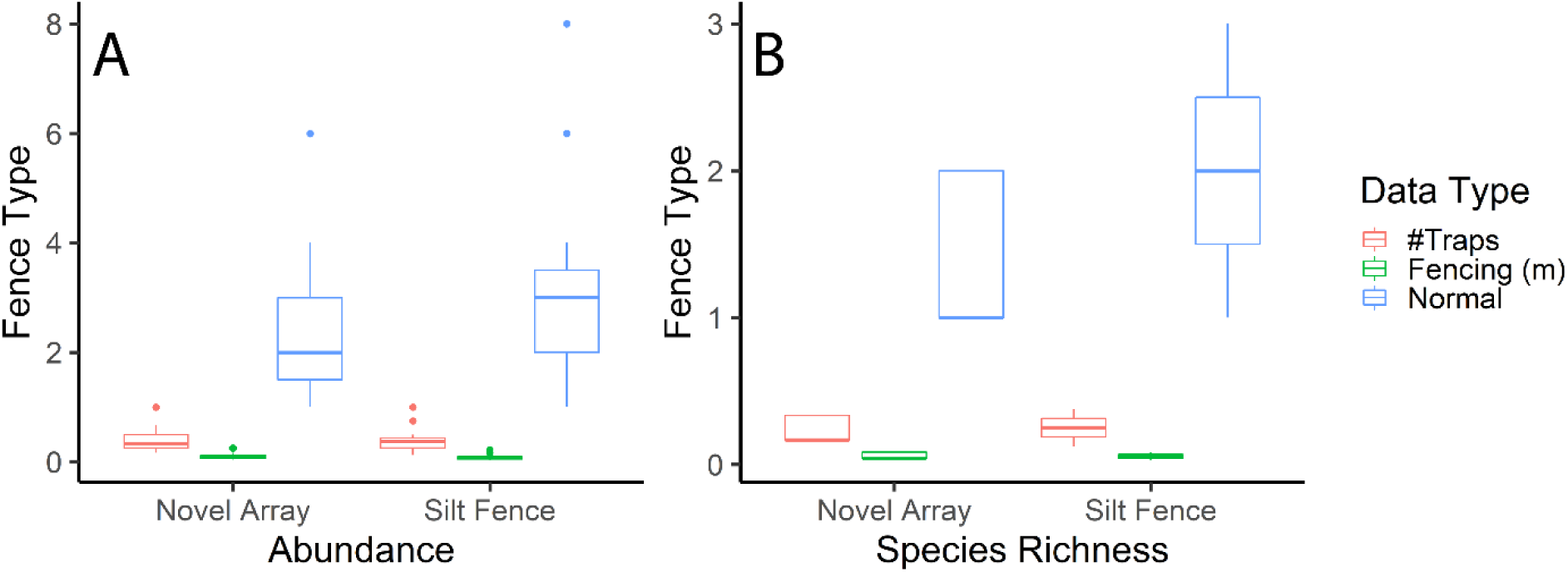
Box and whisker plot demonstrating abundance (A) and species richness (B) for both drift fence types used at the Sevilleta Northeast site. Red indicates that abundance and richness data is controlled for the number of traps used in each drift fence type. Green indicates that abundance and richness data is controlled for the length of fencing in meters. Blue indicates uncorrected abundance data.

**Table 3.** Kruskal-Wallis Test results for both total abundance (’^A^’) and species richness (’^R^’) comparing silt fencing drift fence arrays in 2021 to our novel drift fence array design in 2022 at the Sevilleta Northeast site. ’^T^’ indicates that abundance or richness data was controlled for the number of traps used in each drift fence array type (see Table 2). ‘^M^’ indicates that abundance or richness data was controlled for the length of fencing used in each drift fence array type.

| Test | H-value | P-value |
| --- | --- | --- |
| $n^A$ | 1.1812 | 0.2771 |
| $n^{AT}$ | 0.00439 | 0.9472 |
| $n^{AM}$ | 0.79892 | 0.3714 |
| $n^R$ | 4.0397 | 0.04444 |
| $n^{RT}$ | 0.25614 | 0.6128 |
| $n^{RM}$ | 0.09221 | 0.7614 |

We used PERMANOVA to test for the effect of array design on community composition using the abundance-based Bray-Curtis dissimilarity index [p-value=0.25; F-statistic=1.3477] and the incidence-based Jaccard dissimilarity index [p-value=0.185; F-statistic=1.488]. Again, differences in reptile community composition between the array designs were not statistically significant for either dissimilarity index.

## Discussion

Design options for portable drift fence arrays are rare. This is unfortunate, given the many experimental scenarios where rapid and fast installation, uninstallation, and reinstallation of drift fence arrays are desirable. We present a novel, cost-effective, drift fence array specifically designed for portability in desert conditions. Though primarily developed for rapid deployment and deconstruction, our trapping array design has the added benefit of minimal ground disturbance, which can be a major concern in shallow desert soils with important archaeological sites.

Importantly, there was no statistically significant difference in the capture rate of reptiles when comparing our novel design to the standard silt fencing approach (see Table 3) based on total reptile abundances or reptile community composition. For reptile species richness, our novel array design captured more species, more species per trap and more species per length of fencing over the entire 11 day period (see Table 2). Further, while a Kruskal-Wallis Test (raw species richness values) suggested that total species richness may be greater for the silt fencing design (see ‘Results’& Figure 3), once capture rates were normalized for either trap number or fence length, these differences were not statistically significant (see Table 3). Thus, portability and wind resistance offered by our novel drift fence array design does not appear to come at the expense of capture rate, at least in comparison to silt fence trapping arrays.

The cost of our 24 m drift fence array was $159.39 (see ‘Methods’). This is more expensive than silt fencing alternatives (ca. $23–25, assuming $30 for a 30.48 m [100 ft] roll) (Enge, 1997). However, for study designs with frequent movement between sites, the differences in fencing costs are easily compensated by the reduced labour costs associated with trap portability. Further, while more expensive, PVC posts and bases (contributing to most of the cost discrepancy between our design and standard silt fence arrays) are much more durable than the wooden stakes associated with silt fences. Wooden stakes frequently break upon removal due to user error or natural degradation under chronic weathering in extremely dry or moist conditions. This makes silt fencing even less attractive to redeploy across sites. The portable drift fence array design presented in Rice et al. (2006) costs $47.55 for a fence length of 3.05 m. Extrapolating to the length of fencing in our arrays (24 m), the total cost would be ca. $380.40, which is much higher than our cost of $159.39. Of course, material costs may not scale 1:1 with fence length. Further, the design implemented by Rice et al. (2006) included PVC pipe refugia for treefrogs – an additional feature to terrestrial trapping – that added to the costs of their design. While our design’s PVC posts potentially enable this same feature, testing the utility of PVC as refugia was outside of the purpose of this study.

Even given our trapping success, we made some modifications to our trapping arrays throughout the field season. We originally used cable ties that were 0.508 cm [0.2 in] wide. These frequently broke, either due to sun exposure, transportation, or both. Thus, we eventually implemented a stronger 53.76 kg [120 lb] tensile strength cable tie (length 27.94 cm [11 in] **×** width 0.9 cm [0.354 in]) that alleviated problems with breakage. Further, due to wind stress and/or transportation, the screen fibres occasionally unravelled from the cable ties attached to the PVC posts. This problem was largely solved by folding the screen fabric back onto itself 2–3 times (similar to folding the lower screen edge upon itself before staking to the ground) before cable-tying it to the PVC posts. A duct tape border on its own was not sufficient to prevent screen wear and tear.

We designed fences to be short (8 m long) and used three fences instead of four for enhanced portability. We did so because we were targeting unique whiptail lizard individuals (i.e., we wanted to minimize recapture rates) and thus optimized our design for the ability to move fence arrays both within and between sites in the instance that the rate of unique whiptail captures saturated at any given trapping array. Outside the context of our specific project, however, it may be desirable to modify our general construction framework. This could, for instance, include altering geometries of fences, altering segment lengths, or altering the number of segments per fence. Currently, we are uncertain how the efficacy of our fence materials may differ if implementing these alternative designs. We do not expect that an increase in the number of fences would negatively impact trapping efficacy, especially if folding the fiberglass mesh screens’ vertical edges back on themselves multiple times before piercing the newly created fold with a heavier (wider) cable tie when attaching the mesh screen to the PVC post. However, we suspect that dramatically increasing the number of segments per fence may cause some additional challenges for transportation and wear and tear. Further, we suspect segment length (i.e., the ratio of screen length to fence post frequency) may make the trapping arrays more vulnerable to high wind speeds. In the future, field studies could be used to compare different fence designs with our combination of materials. Further, while our novel drift fence array design appeared successful and provided similar capture rates in a cross-year comparison, a more thorough within-habitat comparative field study should be implemented to better evaluate relative capture rates, abundance, and species richness across array types.

## Acknowledgements

We thank Thomas Dempster, Eva Purcell, and August Spencer for their assistance in the field. Lilly Margeson, Eva Purcell, Simon Dunn, Georgianna Bellinger, Henry Egloff, Kaila Hodges, Camryn Lachica, and Savannah Utz assisted assembling the trapping arrays. This work was completed under the New Mexico Department of Game and Fish permit authorization #3772, the Sevilleta National Wildlife Refuge Special Use Permit #SEV_Bewick_Camper_2022_59, and the USDA-ARS Jornada study permit #592. This study was funded by the NSF award #2105604 and a Clemson University Support for Early Exploration and Development (CUSEED) Grant. All research was approved by Clemson University under IACUC protocol numbers #2020-015 and #2021-047.

---

**Appendix 1.** Abundance values by species for each site surveyed in summer 2022 using our novel drift fence array. Headers indicate the site name (see Table 1). Given their complex taxonomic history and recent taxonomic changes, we used Walker et al. (2021) to reference *Aspidoscelis* species taxonomy and used (B. I. Crother, 2017) to reference other species’ taxonomy. See Table 1 for site name information.

| Taxa | BLM-NW | S-NW | S-NE | J-M | J-MS |
| --- | --- | --- | --- | --- | --- |
| **Snakes** |  |  |  |  |  |
| *Arizona elegans* | - | 1 | - | - | 1 |
| *Coluber flagellum* | - | 1 | 1 | - | 9 |
| *Pituophis catenifer* | - | - | 1 | - | - |
| **Lizards** |  |  |  |  |  |
| *Aspidoscelis inornatus* | - | - | 2 | - | - |
| *Aspidoscelis marmoratus* | 50 | 95 | - | 48 | 143 |
| *Aspidoscelis neomexicanus* | 2 | 17 | 33 | 2 | 9 |
| *Aspidoscelis tessellatus* | 1 | - | - | - | - |
| *Gambelia wislizeni* | - | 1 | - | - | 1 |
| *Holbrookia maculata* | - | - | 1 | - | - |
| *Phrynosoma cornutum* | 9 | 1 | - | - | 5 |
| *Phrynosoma hernandesi* | - | - | 3 | - | - |
| *Phrynosoma modestum* | - | 1 | - | - | - |
| *Plestiodon obsoletus* | - | - | - | - | 2 |
| *Sceloporus bimaculosus* | 1 | 2 | - | - | - |
| *Uta stansburiana* | 3 | 9 | - | 9 | 16 |

